# Atypical hypnotic compound ML297 restores sleep architecture following emotionally-valenced learning, to promote memory consolidation and hippocampal network activation during recall

**DOI:** 10.1101/2022.07.15.500268

**Authors:** Jessy D. Martinez, William P. Brancaleone, Kathryn G. Peterson, Lydia G. Wilson, Sara J. Aton

**Affiliations:** Department of Molecular, Cellular, and Developmental Biology, University of Michigan, Ann Arbor, MI 48109; Undergraduate Program in Neuroscience, University of Michigan, Ann Arbor, MI 48109

## Abstract

Sleep plays a critical role in consolidating many forms of hippocampus-dependent memory. While various classes of hypnotic drugs have been developed in recent years, it remains unknown whether, or how, some of them affect sleep-dependent memory consolidation mechanisms. We find that ML297, a recently-developed candidate hypnotic agent targeting a new mechanism (activating GIRK1-subunit containing G-protein coupled inwardly rectifying potassium [GIRK] channels), alters sleep architecture in mice over the first 6 h following a single-trial learning event. Following contextual fear conditioning (CFC), ML297 reversed post-CFC reductions in NREM sleep spindle power and REM sleep amounts and architecture, renormalizing sleep features to what was observed at baseline, prior to CFC. Renormalization of post-CFC REM sleep latency, REM sleep amounts, and NREM spindle power were all associated with improved contextual fear memory (CFM) consolidation. We find that improvements in CFM consolidation due to ML297 are sleep-dependent, and are associated with increased numbers of highly-activated dentate gyrus (DG), CA1, and CA3 neurons during CFM recall. Together our findings suggest that GIRK1 channel activation restores normal sleep architecture - including REM sleep, which is normally suppressed following CFC - and increases the number of hippocampal neurons incorporated into the CFM engram during memory consolidation.

**Significance Statement:** Both REM and NREM sleep are thought to be important for consolidating hippocampus-dependent memories. We find that GIRK1 activator ML297, administered after single-trial fear learning, restores REM sleep that is normally suppressed after learning fearful associations. This restoration is associated with improvements in fear memory storage, resulting in more robust hippocampus activation in the context of subsequent memory recall. Thus this drug, which also has antiepileptic and anxiolytic properties, may be useful for promoting normal, restorative sleep that benefits memory storage.

## Introduction

Sleep plays an essential role in memory consolidation (1–4). Available data from both human subjects and animal models have implicated both non-rapid eye movement (NREM) and REM sleep in the process of memory storage. While the underlying mechanisms are still under investigation, these states differ one another, and from wake, with respect to neuromodulation, neural network oscillatory behavior, neuronal firing patterns, gene expression, and protein translation (3–13).

Sleep loss over the first few hours following training on many hippocampus-dependent tasks leads to long-term memory disruption (1, 14, 15). Among these, one of the most well-studied is contextual fear memory (CFM), which is initiated by single-trial contextual fear conditioning (CFC) in mice, and is consolidated in a sleep-dependent manner (6, 11, 16–18). Following CFC, both NREM and REM sleep are altered (18–22). Some of these changes, including enhancements in REM theta (4-12 Hz), NREM spindle (7-15 Hz) and NREM sharp wave-ripple oscillations over the first hours following conditioning, predict successful CFM consolidation and recall (18–22). Both CFC and tone-cued fear conditioning also affect sleep architecture in mice, including transiently suppressing REM sleep (23, 24). How REM suppression affects CFM consolidation remains unknown. However, data from analogous studies with human subjects have suggested that post-conditioning REM sleep time, and limbic system brain activation during REM, predicts successful fear memory consolidation (25–27). Moreover, in mice, theta oscillations present in the dorsal hippocampus during post-CFC REM sleep have been shown to play a causal role in promoting CFM consolidation (18, 28). Optogenetically-driven hippocampal theta activity can even rescue CFM consolidation from the deleterious effects of post-CFC sleep deprivation (SD) (18). Thus taken together, available data suggest that limbic system activity and oscillations associated with both NREM and REM sleep contribute to the long-term storage of recently-encoded fear memories.

Hypnotic drug interventions have recently been used as an experimental strategy to test the relationships between sleep, memory consolidation, and synaptic plasticity (29–35). The majority of the hypnotics used in these studies - including benzodiazepines, nonbenzodiazepine “z-drugs”, and sodium oxybate - act as positive allosteric modulators of GABA_A_ receptors or as GABA_B_ receptor agonists. These drugs, while effective at promoting NREM sleep, can have unwanted side effects, including over-sedation, electroencephalogram (EEG) anomalies including aberrant oscillations, and memory deficits (29, 30, 34, 36–39). Recent work has aimed to develop new classes of hypnotic drugs, including orexin receptor antagonists, melatonin receptor agonists, and most recently, activators of G-protein inward rectifying potassium (GIRK) channels (40–42). GIRK channels consist of four subunits (1-4 or K_ir_3.1-3.4), with homo- or hetero-tetrameric compositions that are specific to organs, brain regions, and cell types (e.g. GIRK1/2 channels are selectively expressed in hippocampal neurons, GIRK1/4 channels are present in cardiac myocytes, and GIRK2/3 channels are present in the midbrain) (43–48). GIRK channel activation is typically associated with G_i_-mediated intracellular signaling and itself causes neuronal hyperpolarization, leading to reduced neuronal activity (49, 50). Recent studies have found that the GIRK1 subunit can be directly activated independently of G_i_ using a selective and potent compound known as ML297 (51, 52). Behavioral studies using ML297 in rodents have shown it suppresses seizures, reduces anxiety-like behaviors, and promotes NREM sleep during the circadian active phase (i.e., dark phase) (40, 52, 53). However, it remains unclear whether, and how, ML297 affects sleep-dependent memory processing.

To characterize the effects of GIRK1 channel activation on post-learning sleep and sleep-dependent memory consolidation, we administered ML297 immediately following CFC and measured changes in post-conditioning sleep architecture. We found that while post-CFC NREM sleep was unchanged, ML297 administration restored REM sleep in the hours following CFC (renormalizing it to levels seen at baseline), and significantly improved CFM consolidation. This effect was sleep-dependent - i.e., ML297 had no beneficial effect on CFM when administered during post-CFC SD. Finally, we found that post-CFC sleep, and particularly ML297-augmented post-CFC sleep, led to increased hippocampal cFos and Arc expression during CFM recall. Taken together, our data demonstrate that post-CFC REM sleep plays a critical role in CFM consolidation, leading to greater hippocampal activation during recall - and that restoration of normal post-CFC REM sleep by ML297 promotes this process.

## Materials and Methods

### Animal Handling and Husbandry

All mouse husbandry, experimental, and surgical procedures were reviewed and approved by the University of Michigan Internal Animal Care and Use Committee. For all experiments, 4– 5-month-old, male C57BL/6J mice (Stock No. 000664, Jackson Labs) were housed under a 12:12h light/dark cycle (lights on at 9 AM) and had *ad lib* access to food and water. Mice were housed with littermates until either EEG implantation surgery or (for non-implanted mice) daily habituation prior to behavioral procedures, at which point they were single housed in standard cages with beneficial environmental enrichment.

### Experimental Design and Statistical Analyses

Male littermates were randomly assigned to treatment groups (*n* = 5-7 per group) at the time of single housing for EEG implantation or behavioral procedures. Data analyses were carried out in a blinded manner; in some cases (e.g., for EEG recordings), data were consensus scored by 2 individuals in order to reduce variability. Statistical analyses were carried out using GraphPad Prism software (Version 9.1). For each specific data set, the statistical tests and p-values used are listed in the “Results” section and in corresponding figures and figure legends.

### Surgical Procedures and EEG Recording

For EEG experiments, mice underwent surgical procedures for implantation of electroencephalogram (EEG) and electromyogram (EMG) electrodes. Briefly, mice were anesthetized with 1-2% isoflurane. Stainless steel screw electrodes for EEG recording and referencing were positioned over primary visual cortex (2.9 mm posterior to Bregma, 2.7 mm lateral) bilaterally and cerebellum, respectively, and a braided stainless steel wire EMG electrode was placed in the nuchal muscle. After 11 days of postoperative recovery, each mouse underwent 3 days of habituation to daily handling (5 min/day) and tethering to recording cables in their home cage. Following habituation, 24-h baseline recordings were made from each mouse, starting at lights-on (ZT0). Subsequently, for studies of sleep-dependent memory consolidation, mice underwent CFC training at lights on (ZT0) the following day, and were recorded for an additional 24 h thereafter. EEG/EMG signals (0.5-300 Hz) were amplified at 20 ×, digitized, further digitally amplified at 20-100 ×, and continuously recorded (with a 60-Hz notch filter) using Plexon Omniplex software and hardware (Plexon Inc.) as previously described (18, 19, 54, 55).

### Sleep State and Power Spectra Analysis

Baseline and post-CFC recordings were scored in 10-second epochs as wake, NREM, or REM sleep using custom MATLAB software. EEG and EMG data were band-pass filtered at 0-90 Hz and 150-250 Hz, respectively, for viewing during scoring. Raw EEG data (0.5-300 Hz) were used for fast-Fourier transform and generation of power spectral density from 0.5 to 20 Hz using NeuroExplorer 5 software (Plexon Inc.). An automated spindle detection algorithm was used to identify sleep spindles in band-pass filtered EEG data (7-15 Hz), as intervals containing ≥ 6 successive deviations (i.e., peaks or troughs) of signal that surpassed mean signal amplitude by 1.5 standard deviations, lasting between 0.25-1.75 seconds (55).

### CFC, Drug Administration, Sleep Monitoring, and Sleep Deprivation

Mice underwent single-trial CFC as previously described (6, 11, 18–20). Each mouse was placed in a novel cylindrical conditioning chamber made of clear Plexiglas with a metal grid floor and distal cues (Med Associates). Mice were allowed to freely explore for 2 min and 28 s, after which they received a 0.75 mA, 2-s foot shock through the grid floor, followed by an additional 30 s in the CFC chamber. Immediately following CFC, mice were returned to their home cage and given an i.p. injection of either ML297 (30 mg/kg; Tocris) or vehicle (2% DMSO in 0.5% hydrooxypropyl cellulose aqueous solution). Injections occurred within 5 min of removal from the CFC chamber. CFM tests were conducted 24 h later by returning mice to the CFC chamber for 5 min. Mice were video monitored continuously during both CFC training and CFM testing, and both freezing behavior and locomotor activity (i.e., distance traveled) in the conditioning chamber was quantified in a semi-automated manner using Ethovision XT 16 software (Noldus). CFM-mediated freezing behavior was quantified by subtracting each mouse’s % freezing time during pre-shock baseline from % freezing time across the entire CFM test.

Following CFC, mice were either allowed *ad lib* sleep (Sleep) or were sleep-deprived (SD) via gentle handing over the next 6 h (ZT0-6). Following SD, all mice were allowed *ad lib* recovery sleep over the next 18 h prior to CFM testing. For mice without EEG/EMG implants, sleep was quantified over the first 6 h post-CFC via visual monitoring. Every 5 min, individual mice were scored as awake or asleep, with sleep identification based on immobility, slow breathing, and presence of stereotyped (crouched) sleep postures, consistent with prior studies. For SD, gentle handling procedures included cage tapping or shaking, or nest disturbance (Fisher et al., 2012; Delorme et al., 2019, 2021; Puentes-Mestril et al., 2021).

### Histology and Immunohistochemistry

To quantify hippocampal activation patterns associated with recall, 90 min following the conclusion of CFM tests, mice were euthanized with an overdose of sodium pentobarbital and perfused with ice cold PBS, followed by ice cold 4% paraformaldehyde. Brains were dissected, post-fixed, and cryoprotected in a 30% sucrose solution. 50 µm coronal dorsal hippocampal sections were immunostained using rabbit-anti-cFos (1:1000; Abcam, ab190289) and guinea pig-anti-Arc (1:500; Synaptic Systems, 156004) as markers of neuronal activation. Secondary antibodies used included Alexa Fluor 488 (1:200; Invitrogen, A11032) and Alexa Fluor 594 (1:200; Invitrogen, A11034). Stained sections were mounted using Prolong Gold antifade reagent with DAPI (Invitrogen, P36931) and imaged using a Leica SP8 confocal microscope with a 10X objective, to obtain z-stack images (10 µm steps) for maximum projection of fluorescence signals. Identical image acquisition settings (e.g. exposure times, frame average, pixel size) were used for all sections.

For analysis of hippocampal activation patterns, three images of dorsal hippocampus were taken per mouse and equally sized regions of interest (ROIs) for DG, CA1 and CA3 regions were obtained for each image. cFos+ and Arc+ neurons were identified and quantified in subregions of these ROIs (i.e., pyramidal or granule cell layers, DG hilus) by a scorer blinded to animal condition, using ImageJ software. For Arc expression in pyramidal cell layers of CA1 and CA3, mean fluorescent intensity (MFI) was measured by adaptive thresholding of fluorescent signals and subtracting the fluorescence intensity of each region from mean background fluorescence (12).

## Results

### GIRK channel activation renormalizes REM sleep architecture following CFC and improves CFM consolidation

To test how GIRK1 channel activation affects post-CFC sleep and sleep-dependent CFM consolidation, we recorded ML297-induced changes in sleep architecture and EEG activity following CFC (**Fig. 1–2**). 4-5 month old, male C57BL/6J mice underwent continuous 24-h baseline EEG/EMG recording starting at lights-on (ZT0), followed by single-trial CFC at ZT0 the following day. Immediately following CFC, mice were administered either vehicle or ML297 (30 mg/kg, i.p.), and underwent EEG recording for additional 24 h prior to CFM testing at ZT0 on the third day (**Fig. 1A**). Semi-automated scoring during CFC and CFM testing was used to quantify both locomotor activity (i.e., travel distance) within the CFC chamber and CFM-associated freezing behavior (**Fig. 1B-C**). During initial training (prior to foot shock), vehicle- and ML297-treated mice showed similar locomotor activity in the CFC chamber (two-tailed, unpaired t-test; *p* = 0.5622; t, df = 0.5994, 10) and comparable, low amounts of freezing (two-tailed, unpaired t-test; *p* = 0.2663; t, df = 1.177, 10) (**Fig. S1**). However, during CFM testing, locomotion was reduced (two-tailed, unpaired t-test; *p* = 0.0392; t, df = 2.372, 10) and freezing was enhanced (two-tailed, unpaired t-test; *p* = 0.0101; t, df = 3.162, 10) in ML297-treated mice (compared to vehicle-treated controls; **Fig. 1C**). This suggests that sleep-dependent CFM consolidation is improved by post-CFC administration of ML297.

**Figure 1:**
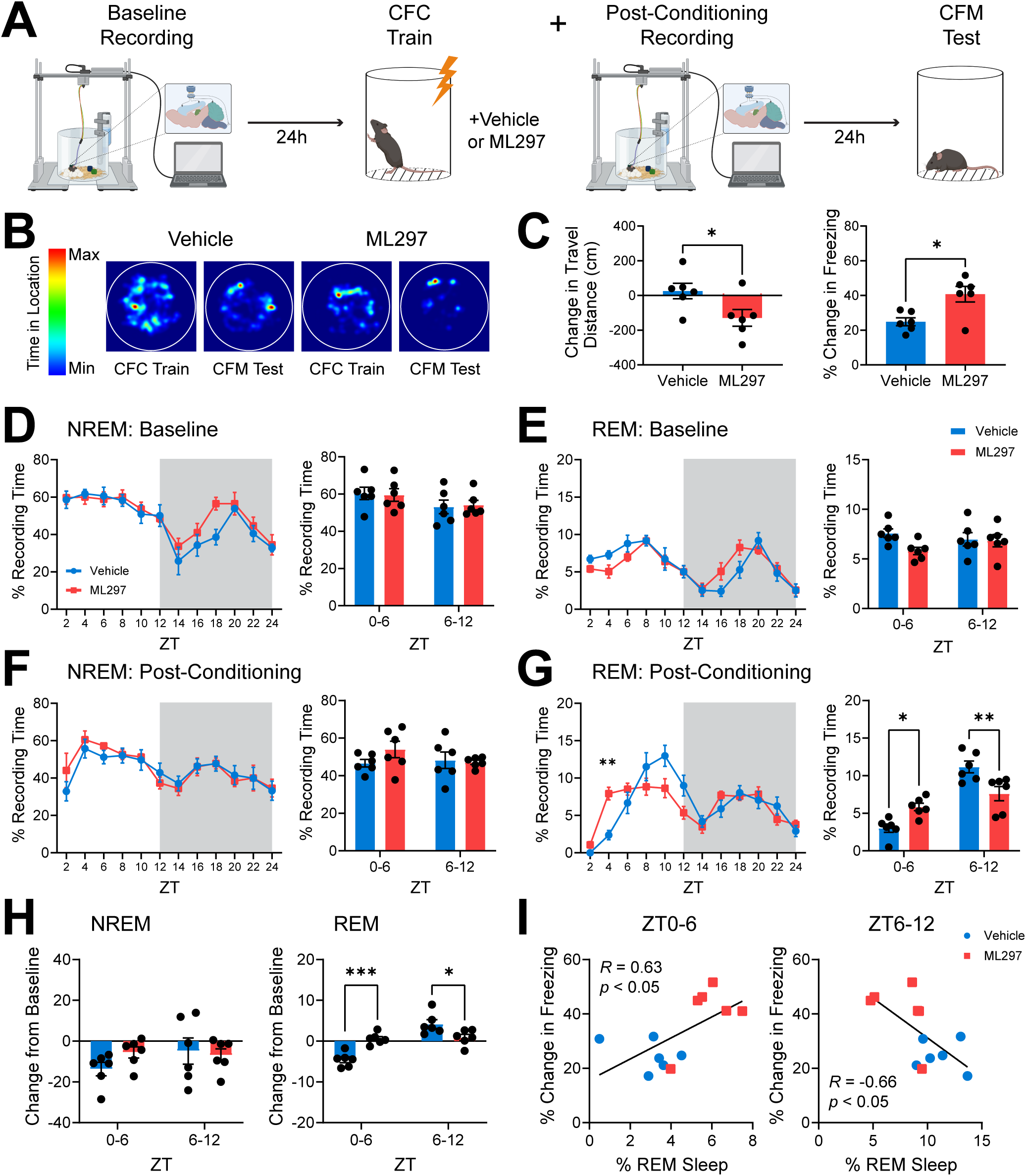
GIRK1 activation renormalizes REM sleep amounts during fear memory consolidation and improves fear memory recall. (***A***) Configuration of EEG electrodes for sleep recording and schematic of experimental design. Mice were recorded over a 24-h baseline starting at lights-on (ZT0), then underwent single-trial contextual fear conditioning (CFC) followed by an i.p. injection of vehicle or GIRK1 activator ML297 (30 mg/kg). Each mouse was recorded for an additional 24 h prior to being tested for contextual fear memory (CFM). (***B***) Heat maps of time in various locations within the conditioning chamber, for representative vehicle- and ML297 treated mice during CFC training and CFM testing. (***C***) During CFM testing, the reduction in travel distance was significantly lower, while increases in freezing behavior (from pre-shock values during prior conditioning) were significantly higher, in ML297-treated mice. Bars indicate mean ± SEM; *n* = 6 mice/group; * indicates *p* = 0.0392 (change in travel distance), * indicates *p* = 0.0101 (change in freezing), two-tailed, unpaired t-test. (***D***) NREM and (***E***) REM sleep behavior during baseline across the light:dark cycle for vehicle- and ML297-treated mice. Gray shaded areas represent lights off. No changes seen in time spent in NREM or REM sleep across the light:dark cycle nor in total NREM or REM sleep across 6 h quartiles. *n* = 6 mice/group (***F-G***) Sleep behavior post-conditioning across the light:dark cycle for vehicle- and ML297-treated mice. Gray shaded areas represent lights off. No changes seen in time spent in NREM across the light:dark cycle. Time spent in REM sleep was significantly altered during the light cycle. REM sleep is significantly increased 3-4 hours post-treatment of ML297. Values indicate mean ± SEM; *n* = 6 mice/group; ** indicates *p* = 0.0057, Sidak’s *post hoc* test vs. vehicle. ML297-treated also had significant more total REM sleep during ZT0-6 and significantly less total REM sleep during ZT6-12. Sidak’s *post hoc* test vs. vehicle. * indicates *p* = 0.0226 (ZT0-6) and *p* = 0.0045 (ZT6-12), Sidak’s *post hoc* test vs. vehicle. (***H***) Compared to baseline, REM sleep in the first 6 h post-CFC was more suppressed in vehicle-treated vs. ML297-treated mice, while significantly promoted during the last 6 h post-CFC. *** and * indicates *p* = 0.0004 (ZT0-6) and *p* = 0.0169 (ZT6-12), respectively, Sidak’s *post hoc* test vs. vehicle. (***I***) Correlation between freezing behavior and % time spent in REM sleep in the first 6 h post CFC and last 6 h post CFC of the light cycle. *R* and *p* values are shown for Pearson correlation.

**Figure 2:**
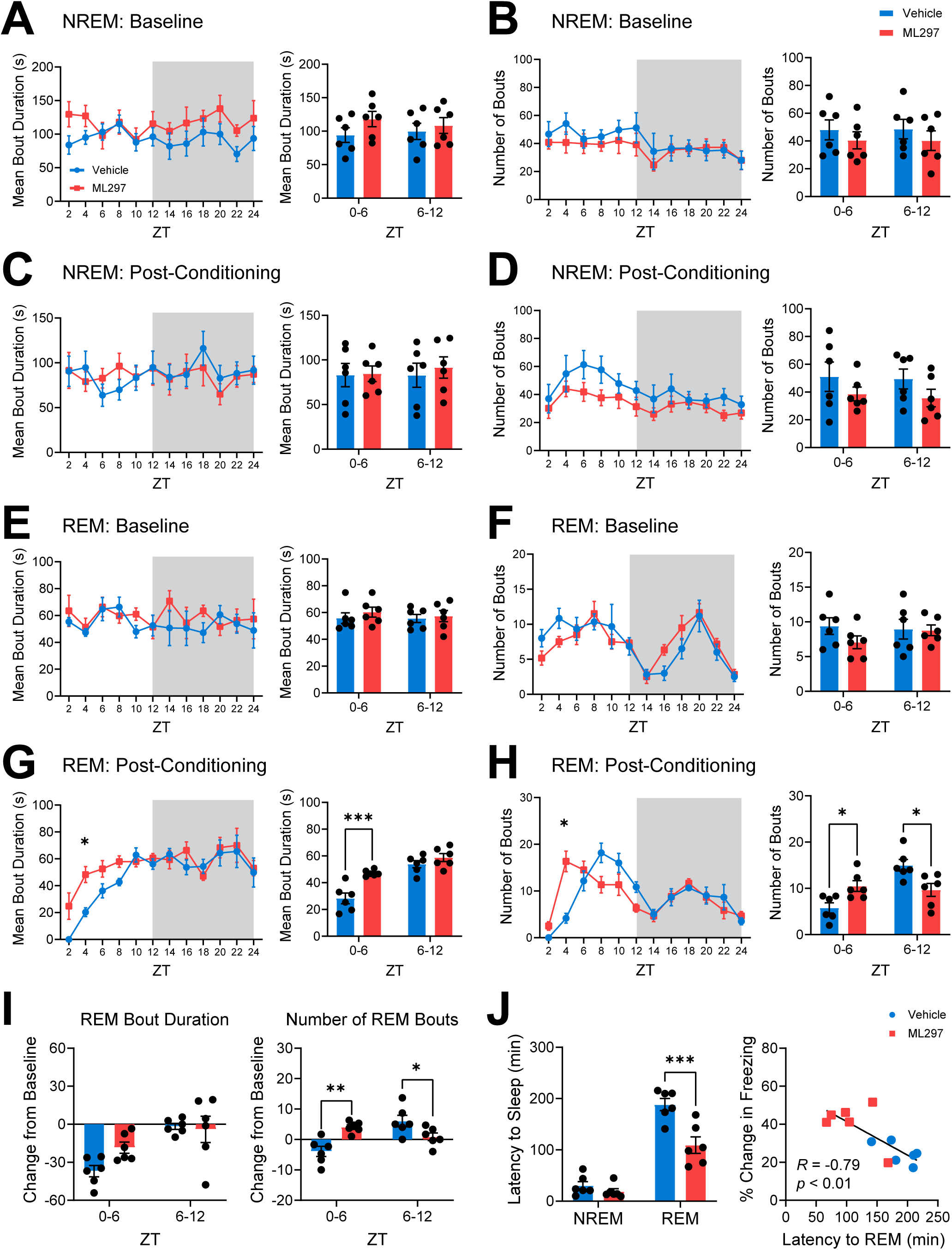
ML297 REM sleep architecture is promoted during post-conditioning with GIRK channel activation. (***A***) Mean bout duration and (***B***) number of bouts for NREM sleep over the 24 h during baseline in vehicle- and ML297-treated mice were similar. (***C***) Mean bout duration and (***D***) number of bouts for NREM sleep over the 24 h post-CFC in vehicle- and ML297-treated mice were similar. (***E***) Mean bout duration and (***F***) bout numbers for REM sleep over the 24 h during baseline in vehicle- and ML297 treated mice were similar. (***G***) Mean bout duration and (***H***) bout numbers for REM sleep over the 24 h post-CFC in vehicle- and ML297 treated mice. * indicates *p* = 0.0400 (REM bout duration), *p* = 0.0177 (number of REM bouts), Sidak’s *post hoc* test vs. vehicle. Values indicate mean ± SEM. (***G***) REM mean bout durations and (***H***) bout numbers are shown for hours 0-6 and 6-12 post-CFC. ML297-treated mice show longer REM bout durations during ZT0-6. ** indicates *p* = 0.0003, Sidak’s *post hoc* test vs. vehicle. ML297-treated mice show a greater number of REM sleep bouts during ZT0-6 and a reduced number of REM sleep bouts during ZT6-12. * indicates *p* = 0.0267 (ZT0-6), *p* = 0.0123 (ZT6-12), Sidak’s *post hoc* test vs. vehicle. (***I***) REM mean bout duration and number shown as a change from time-matched baseline values. No changes from baseline were observed in REM bout duration. There is less reduction of REM sleep bouts from baseline over the first 6 h post-CFC period and last 6 h period of the light cycle. ** indicates *p* = 0.0019 (ZT0-6) and * indicates *p* = 0.0419 (ZT6-12), Sidak’s *post hoc* test vs. vehicle. (***J***) ML297-treated mice showed decreased latency to REM sleep (but unchanged latency to NREM sleep) after CFC (*** indicates *p* = 0.0001, Sidak’s *post hoc* test vs. vehicle). Shorter REM latency predicted successful CFM recall testing. *R* and *p* values are shown for Pearson correlation. *n* = 6 mice/group

To test whether enhanced CFM consolidation following ML297 administration correlated with differences in sleep architecture, we first compared baseline vs. post-CFC sleep amounts between vehicle- and ML297-treated mice. Vehicle- and ML297-treated mice showed similar NREM and REM amounts across the 24-hour baseline recording period (**Fig. 1D-E**), as well as similar NREM sleep amounts over the next 24 h post-CFC (two-way repeated measures (RM) ANOVA; *p*(time of day x treatment) = 0.8121; *F*(11, 110 = 0.6157); **Fig. 1F**). However, ML297 significantly increased REM sleep amounts over the first 6 h following CFC (two-way RM ANOVA; *p*(time of day x treatment) = <0.0001; *F*(11,110 = 4.209); Sidak’s *post hoc* test for vehicle vs. ML297, *p* = 0.0226; **Fig. 1G**), and reduced REM (relative to vehicle-treated controls) over the following 6 h (ZT6-12; Sidak’s *post hoc* test for vehicle vs. ML297, *p* = 0.0045). While CFC reduced REM sleep overall relative to baseline in vehicle-treated mice (two-way RM ANOVA; *p*(time of day x treatment) = 0.0003; *F*(1,10 = 30.57)), but ML297 reversed this effect over ZT0-6 (Sidak’s *post hoc* test for vehicle vs. ML297, *p* = 0.0004; **Fig. 1E,G-H**). ML297 did not affect the proportion of time spent in NREM at any timepoint compared with vehicle (two-way RM ANOVA; *p*(time of day x treatment) = 0.1657; *F*(1,10 = 2.236)), the change in NREM amounts from baseline following CFC (two-way RM ANOVA; *p*(time of day x treatment) = 0.0749; *F*(1,10 = 3.953); **Fig. 1D, F, H**), or the proportion of time spent in NREM or REM during the dark phase following CFC (ZT12-24) (**Fig. S2**). Critically, freezing behavior at recall was positively correlated with the proportion of time spent in REM sleep over the first 6 h post-CFC (Pearson correlation coefficient, *R* = 0.6319; *p* = 0.0275), but negatively correlated with REM amounts over the subsequent 6 h (i.e., ZT6-12; Pearson correlation coefficient, *R* = −0.6550, *p* = 0.0208; **Fig. 1I**).

We also quantified how CFC and ML297 affected other features of sleep architecture, including NREM and REM bout durations and bout numbers. These aspects of NREM and REM sleep were similar at baseline between vehicle- and ML297-treated mice (**Fig. 2A-B, E-F**). Following CFC, ML297 had no effect on either NREM bout duration (two-way RM ANOVA; *p*(time of day x treatment) = 0.4974; *F*(11,110 = 0.9487); **Fig. 2C**) or bout number (two-way RM ANOVA; *p*(time of day x treatment) = 0.6252; *F*(11,110 = 0.8149); **Fig. 2D**). In contrast, over the first 6 h post-CFC, ML297 significantly increased REM sleep bout durations (two-way RM ANOVA; *p*(time of day x treatment) = 0.0245; *F*(1,10 = 6.994); Sidak’s *post hoc* test for vehicle vs. ML297, *p* = 0.0003; **Fig. 2G**) and bout numbers (two-way RM ANOVA; *p*(time of day x treatment) = 0.0016; *F*(1,10 = 18.39); Sidak’s *post hoc* test, *p* = 0.0267; **Fig. 2H**). While in vehicle-treated mice, CFC reduced both REM bout duration and number (relative to baseline), ML297 restored REM bout numbers to baseline levels (Sidak’s *post hoc* test for vehicle vs. ML297, *p* = 0.0019; **Fig. 2I**). Finally, latency to the first bout of post-CFC REM sleep (but not NREM sleep) was significantly reduced in mice administered ML297 (two-way RM ANOVA; *p*(sleep state x treatment) = 0.0052; *F*(1,10 = 12.67); Sidak’s *post hoc* test for vehicle vs. ML297, *p* = 0.0001; **Fig. 2J**). Reduced latency to REM following CFC also predicted successful CFM recall the following day, with mice with the shortest latency to REM showing the highest levels of freezing (Pearson correlation coefficient, *R* = −0.7924, *p* = 0.0021; **Fig. 2J**).

Taken together, these data suggest that ML297-mediated GIRK channel activation may improve CFM consolidation through a restorative increase in REM sleep over the first few hours following CFC. Thus, ML297 treatment has the effect of renormalizing REM sleep architecture, to offset suppression of REM that typically occurs after CFC and other fear-associated learning (56).

### ML297 administration alters NREM and REM EEG oscillations in a manner consistent with renormalizing sleep architecture

We next assessed how NREM- and REM-associated EEG oscillations are affected by ML297 administration. We found that at baseline, vehicle- and ML297-treated mice showed no EEG spectral power differences in either NREM or REM sleep (**Fig. 3A, C**). Following CFC, vehicle- and ML297-treated mice showed NREM spectral power differences (two-way RM ANOVA; *p*(frequency x treatment) < 0.0001; *F*(78, 1343 = 3.643)) with vehicle-treated mice having a greater proportion of total EEG power in the NREM delta (0.5-4 Hz) band (**Fig. 3B**). Post-CFC REM EEG spectra also differed between groups, with ML297-treated mice having greater proportional spectral power in the theta (4-12 Hz) band (two-way RM ANOVA; *p*(frequency x treatment) < 0.0001; *F*(78, 869 = 2.349), Sidak’s *post hoc* test for vehicle vs. ML297, *p* = 0.001; **Fig. 3D**).

**Figure 3:**
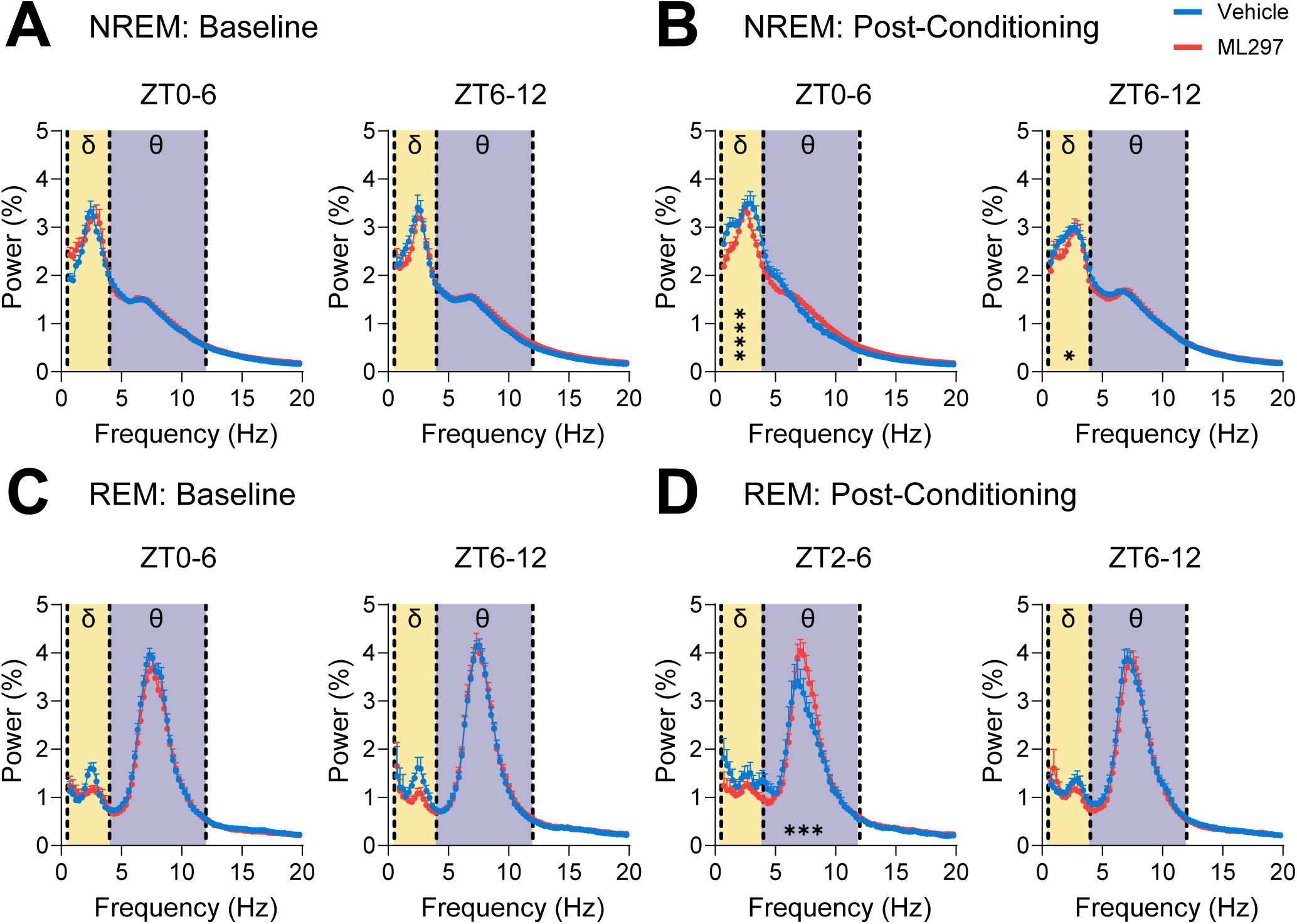
ML297 has modest effects on overall NREM and REM EEG spectral power. EEG power spectra (recorded over visual cortex, bilaterally) are shown for vehicle- and ML297-treated mice during NREM baseline (***A***) and post-CFC (***B***), and during REM baseline (***C***) and post-CFC (***D***). Values indicate % of total spectral power at each frequency band, mean ± SEM; *n* = 6 mice/group. For (***B***), **** and * indicate post-CFC differences in NREM delta frequency bands at ZT0-6 and 6-12, respectively, *p* ≤ 0.0001 for 2.9-3.9 Hz and *p* < 0.05 for 1.4-1.7 Hz, respectively, Sidak’s *post hoc* test vs. vehicle. For (***D***), *** indicates post-CFC differences in REM at ZT2-6, *p* < 0.001 for 7.1-8.1 Hz, Sidak’s *post hoc* test vs. vehicle. Values indicate mean ± SEM; *n* = 6 mice/group.

Because ML297 enhanced post-CFC REM sleep, and caused relative decrease in NREM delta power, we also assessed the effects of both CFC and ML297 on NREM sleep spindles - waxing-and-waning, discrete EEG oscillations with a peak frequency of 7-15 Hz. Spindles are: 1) inversely related to NREM delta power (57–59), 2) implicated in CFM consolidation (21, 60), and 3) critical for transitions between NREM and REM sleep (61, 62). To test whether alterations in REM sleep architecture after ML297 administration were associated with changes in NREM spindles, we next detected these events in a semi-automated manner (55) and compared post-CFC spindle characteristics between groups. Both at baseline, and following CFC, neither spindle density (two-way RM ANOVA; *p*(time of day x treatment) = 0.7795; *F*(5, 50 = 0.4935)) nor duration (two-way RM ANOVA; *p*(time of day x treatment) = 0.7784; *F*(5, 50 = 0.4951)) differed significantly between ML297- and vehicle-treated mice (**Fig. 4A-B, D-E**). Spindle power during baseline NREM sleep was similar between the two groups and relatively invariant across the entire light phase (ZT0-12; **Fig. 4C**). However, the proportion of total NREM EEG spectral power in the spindle (i.e., sigma) band was significantly higher following CFC in ML297-treated mice (two-way RM ANOVA; *p*(time of day x treatment) = 0.0211; *F*(5, 50 = 2.938); **Fig. 4E**). This difference appeared to reflect suppressed spindle power (relative to baseline) in vehicle-treated mice over the first 4 h of NREM following CFC, which were reversed by ML297 (Sidak’s *post hoc* test for vehicle vs. ML297 at ZT0-2 and 2-4, *p* = 0.0223 and *p* = 0.0424). This difference in spindle power between the groups following CFC is consistent with both the relative increases in NREM EEG delta power, and the overall suppression of REM, following conditioning in vehicle-treated (but not ML297-treated) mice. Together, our EEG data suggest that administration of GIRK channel activator ML297 following CFC modestly augments REM theta, and renormalizes NREM delta/spindle ratios, in a manner consistent with its renormalization of sleep architecture. Because these EEG oscillatory changes are similar to those known to be associated with successful sleep-dependent CFM consolidation (18, 19, 21), they are consistent with state-dependent hippocampal oscillations serving as a potential driver of memory enhancement by ML297 (3, 4).

**Figure 4:**
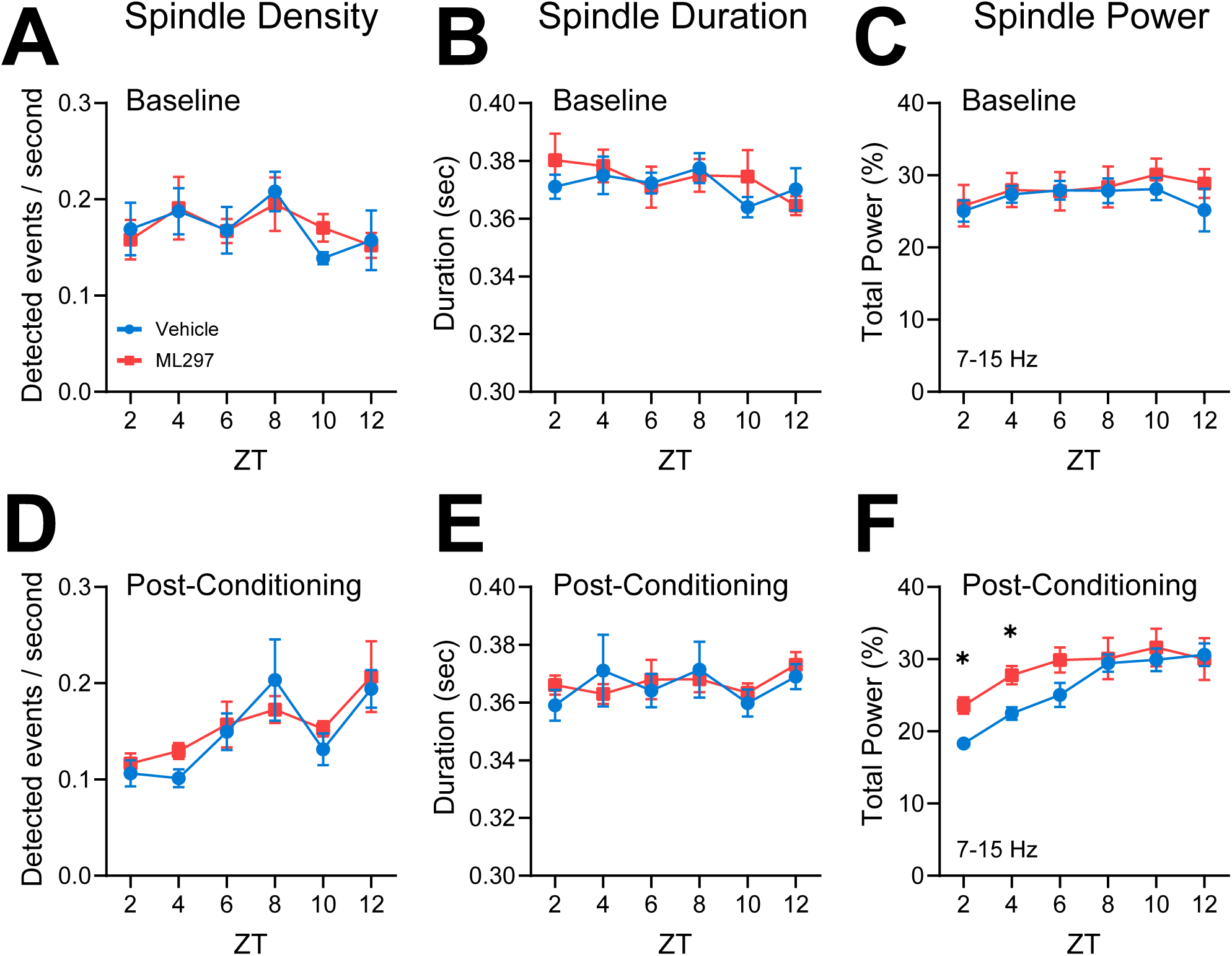
ML297 normalizes NREM sleep spindle power in the hours following CFC. NREM spindle density (***A***) and spindle duration (***B***) were similar between vehicle- and ML297-treated groups for hours 0-12 during baseline. (***C***) NREM EEG spectral power within the spindle/sigma frequency band (7-15 Hz) was similar between groups at baseline. NREM spindle density (***D***) and spindle duration (***E***) were similar between both vehicle- and ML297-treated mice following CFC. (***F***) Average NREM spindle power was reduced in vehicle-treated mice relative to ML297-treated mice, over the first 4 h following CFC * indicates *p* < 0.05, Sidak’s *post hoc* test. Values indicate mean ± SEM; *n* = 6 mice/group.

### ML297 effects on CFM consolidation are sleep-dependent

Because ML297 restores REM sleep architecture and NREM oscillations during CFM consolidation, we next tested whether ML297-mediated improvement in CFM was sleep-dependent, or due to other effects of GIRK activation. In a second cohort of non-instrumented mice, we tested whether post-CFC SD interfered with ML297-driven improvements in CFM consolidation. At lights-on, mice underwent single-trial CFC training and were immediately administered either vehicle or ML297. Over the next 6 h, mice in each treatment group were either allowed *ad lib* sleep (and visually monitored for changes in sleep amount) or underwent gentle-handling SD in their home cage (which is sufficient to disrupt CFM consolidation) (6, 18). CFM recall was tested for all mice 24 h after training (**Fig. 5A**). The total time spent asleep over the first 6 h following CFC was similar for the two freely-sleeping groups, with no significant effect of ML297 on total sleep time (two-tailed, unpaired t-test; *p* = 0.4077; t, df = 0.8684, 9) (**Fig. 5B**). Mice were video monitored during both CFC and CFM recall testing. As expected, during initial CFC training (**Fig. 5C**), locomotor activity and freezing were high and low, respectively, and were similar across treatment groups (**Fig. S4**). 24 h after CFC, mice were returned to the CFC chamber to test CFM recall. During recall, there was a significant effect of both prior sleep condition and drug treatment on locomotor activity (two-way ANOVA; *p*(treatment) = 0.0074, *F*(1, 20 = 8.894); *p*(sleep condition) = 0.0057, *F*(1, 20 = 9.592)) *p*(treatment x sleep condition interaction) = 0.0395, *F*(1, 20 = 4.854)) and freezing (two-way ANOVA; *p*(treatment) = 0.0022; *F*(1, 20 = 12.29); *p*(sleep condition) < 0.0001; *F*(1, 20 = 31.07)). As expected, freely-sleeping vehicle-treated mice had superior CFM comparted to vehicle-treated SD mice (Tukey’s *post hoc* test, *p* = 0.0140) (**Fig. 5D**). Freely-sleeping ML297-treated mice showed stronger CFM than all other groups tested (Tukey’s *post hoc* test vs. Sleep+ML297: Sleep+Vehicle, *p* = 0.0423, SD+Vehicle, *p* < 0.0001, and SD+ML297, *p* = 0.0012). However, SD disrupted CFM consolidation regardless of ML297 treatment - i.e., there was no effect of ML297 when mice were sleep deprived (Tukey’s *post hoc* test for vehicle vs. ML297 in the SD condition, *NS*). These findings support the conclusion that sleep is required for ML297-mediated improvements in CFM consolidation. Together, these data suggest that GIRK1 channel activation promotes memory consolidation via sleep-dependent mechanisms.

**Figure 5:**
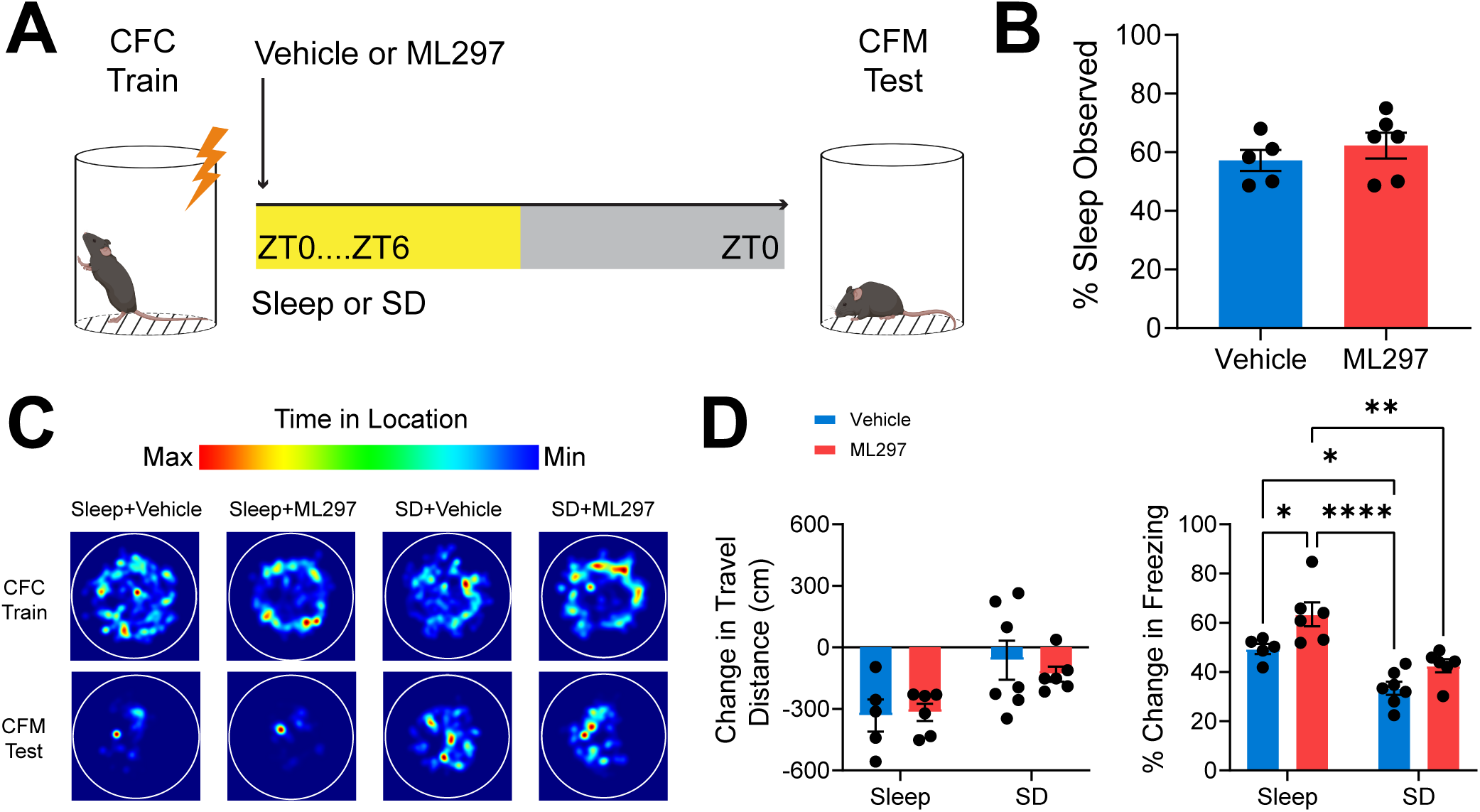
Post-CFC ML297 administration improves CFM consolidation in a sleep-dependent manner. (***A***) Experimental design. Mice underwent single-trial CFC at lights on, were subsequentially administered vehicle or ML297 (30 mg/kg) via i.p. injection and returned to their home cage. Mice in the two drug treatment groups were then allowed *ad lib* sleep or were sleep deprived over the first 6 h post-CFC, after which all mice were allowed to sleep freely in their home cage until CFM testing the following day at lights on. (***B***) Total amounts of observed sleep during the first 6 h post-CFC were similar for the two *ad lib* sleep groups. Bars indicate mean ± SEM, *n* = 5 and 6 mice, respectively, for vehicle and ML297. (***C***) Heat maps of time in various locations within the conditioning chamber, for representative vehicle- and ML297 treated mice during CFC training and CFM testing. (***D***) During CFM, travel distance tended to be more reduced (compared with pre-shock baseline during CFC training) in freely-sleeping mice. Freezing behavior was significantly greater in freely-sleeping ML297-treated mice compared with freely-sleeping vehicle-treated mice. SD reduced CFM-associated freezing behavior in both treatment groups, which did not differ from one another. *n* = 5 and 6 mice, respectively, for vehicle and ML297 with *ad lib* sleep, and *n* = 7 and 6 mice, respectively, for vehicle and ML297 with SD. *, **, and **** indicate *p* < 0.05, *p* < 0.01, and *p* < 0.0001, respectively, Tukey’s *post hoc* test.

### ML297-mediated improvement in CFM consolidation is associated with greater hippocampal activation during subsequent recall

The major input to hippocampus from the neocortex is relayed through the DG. Acting as a gateway to the rest of the hippocampus, the DG receives sensory and non-sensory information from the rest of the neocortex via entorhinal cortical input. Neuronal immediate early gene (IEG) expression increases among DG granule cells during both initial learning and memory retrieval, and granule cell activation plays a causal role in recall (63). We tested whether changes in DG activity during CFM recall were associated with sleep- and ML297-mediated improvements in CFM consolidation, by quantifying cFos and Arc expression in hippocampus. After CFM recall, mice were returned to their home cages; 90 min later they were perfused to quantify protein products of IEG expression associated with recall. We found a significant effects of both sleep and ML297 treatment on cFos expression in the DG (two-way ANOVA; *p*(sleep condition) < 0.0001, *F* (1, 16) = 57.37; *p*(treatment) = 0.0003, *F*(1, 16 = 20.80); **Fig. 6A-B**). ML297-treated mice allowed *ad lib* sleep had significantly increased cFos+ cell counts in DG during recall compared to both SD groups, and freely-sleeping vehicle-treated counterparts (Sidak’s *post hoc* test vs. Sleep+ML297: Sleep+Vehicle, *p* = 0.0116; SD+Vehicle, *p* < 0.0001; SD+ML297, *p* = 0.0002; **Fig. 6B-C**). SD also disrupted recall-associated DG cFos expression in vehicle-treated mice (Sidak’s *post hoc* test, *p* = 0.0010; **Fig. 6C**). Similar recall-associated patterns of were observed for DG Arc expression (two-way ANOVA; *p*(sleep condition) = 0.0009; *F*(1, 16 = 16.49); *p*(treatment) = 0.0304; *F*(1, 16 = 5.641)) (**Fig. 6B,D**), with reduced numbers of Arc+ neurons in SD mice (Sidak’s *post hoc* test vs. Sleep+ML297: SD+Vehicle, *p* = 0.0020; SD+ML297, *p* = 0.0057) (**Fig. 6D**). Overall expression of both IEGs in DG at recall was predictive of successful recall, with higher numbers of cFos+ and Arc+ neurons corresponding to increased freezing behavior during CFM testing (cFos: Pearson correlation coefficient, *R* = 0.7225, *p* = 0.0003; Arc: Pearson correlation coefficient, *R* = 0.5521, *p* = 0.0116) (**Fig. 6E**).

**Figure 6:**
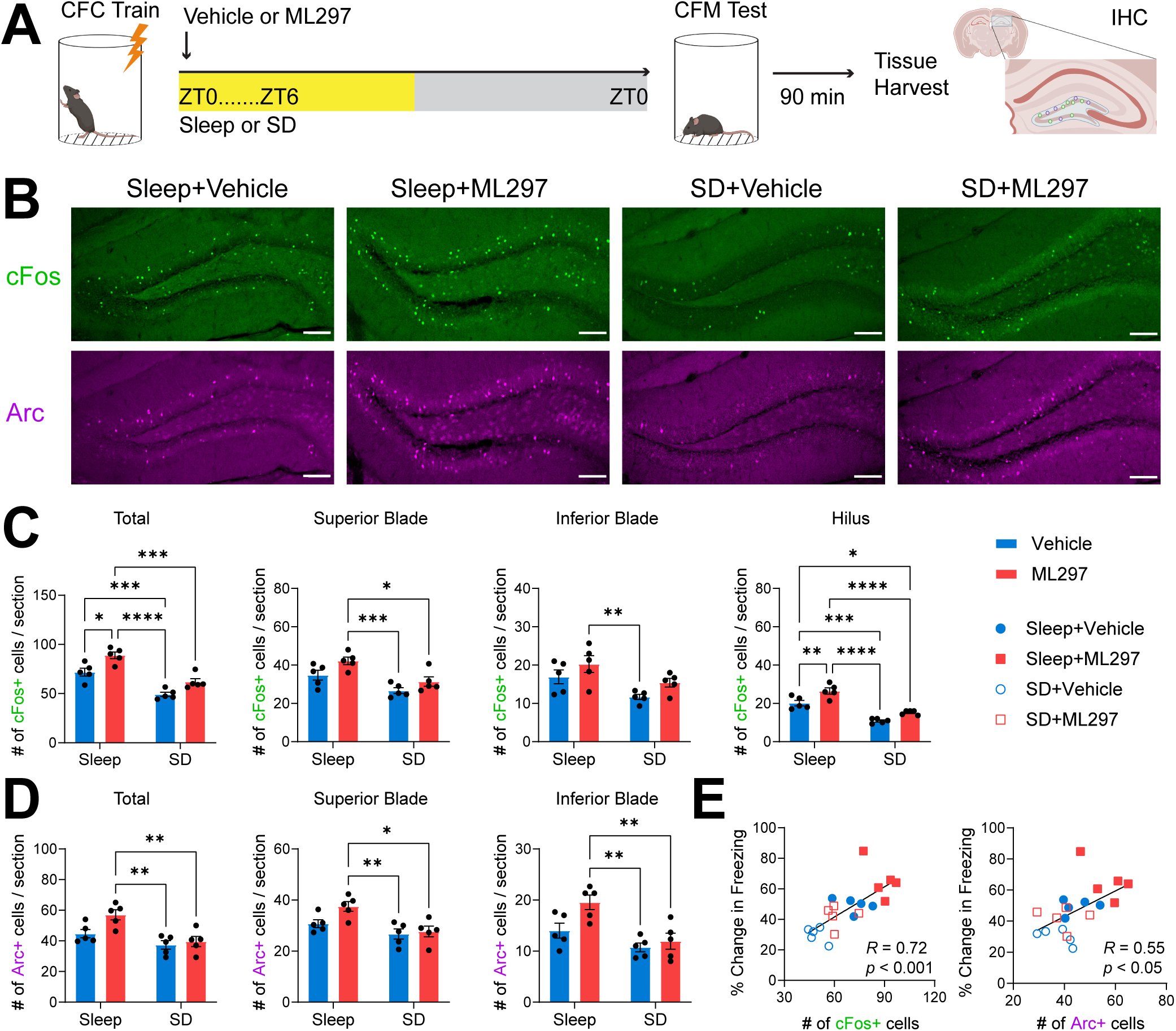
Post-CFC ML297 increases the number of active neurons in DG during subsequent CFM recall, in a sleep-dependent manner. (***A***) Experimental paradigm. Mice underwent single-trial CFC, were administered vehicle or ML297 (30 mg/kg) following training, and then were either allowed *ad lib* sleep or underwent 6-h SD. 24 h after CFC, mice were tested for CFM, and perfused 90 min later to immunohistochemically quantify recall-associated cFos and Arc IEG expression in dorsal hippocampus. *n* = 5 mice/ group. (***B***) Representative images of cFos+ (green) and Arc+ (magenta) DG neurons following CFM recall in the four treatment groups. Scale bar = 100 µm. (***C***) Mice allowed *ad lib* sleep had significantly increased cFos+ neuron counts across DG compared to both SD groups. ML297 increased cFos+ neuron numbers further in freely-sleeping mice. Similar patterns were observed for the two DG granule cell blades, and for the DG hilus. (***D***) Arc+ neuron counts across DG were also reduced in both SD groups. ML297 administration led to a trend for higher overall Arc+ DG neurons relative to vehicle in freely-sleeping mice. *, **, ***, and **** indicate *p* < 0.05, *p* < 0.01, *p* < 0.001, and *p* < 0.0001, respectively, Sidak’s *post hoc* test. (***E***) Higher numbers of cFos+ and Arc+ neurons across DG at recall reflected the success of CFM consolidation across individual mice. *R* and *p* values are shown for Pearson correlation.

We also quantified IEG expression within individual subregions of the DG to examine whether changes associated with recall in the four treatment groups were region-specific. As shown in **Fig. 6C-D**, similar patterns of expression were observed in the granule cell body layer of both the superior and inferior blade of DG, and in the DG hilus. For cFos expression, we found significant effect of both sleep and treatment in both the superior and inferior blades (superior: two-way ANOVA; *p*(sleep condition) = 0.0005, *F* (1, 16) = 19.15; *p*(treatment) = 0.0127 *F*(1, 16 = 7.863); inferior: two-way ANOVA; *p*(sleep condition) = 0.0051, *F* (1, 16) = 10.53; *p*(treatment) = 0.0379; *F*(1, 16 = 5.123)), as well as in the hilus (two-way ANOVA; *p*(sleep condition) < 0.0001, *F* (1, 16) = 82.24; *p*(treatment) = 0.0002; *F*(1, 16 = 22.19); **Fig. 6C**). Similarly, Arc expression patterns in superior and inferior blades after recall were similar to overall Arc+ neuron numbers (superior: two-way ANOVA; *p*(sleep condition) = 0.0021, *F* (1, 16) = 13.37; *p*(treatment) = 0.0584 *F*(1, 16 = 4.155); inferior: two-way ANOVA; *p*(sleep condition) = 0.0010, *F* (1, 16) = 16.03; *p*(treatment) = 0.0246; *F*(1, 16 = 6.154); **Fig. 6D**).

To better understand how recall-associated neuronal activation is affected across the rest of the hippocampal circuit as a function of post-learning sleep and ML297, we also examined IEG expression within the pyramidal cell layers of CA1 and CA3 after CFM recall (**Fig.7A-B**). Recall-driven cFos+ neuron numbers in CA1 varied significantly as a function of prior sleep and drug treatment in CA1 (CA1: two-way ANOVA; *p*(sleep condition) = 0.0060, *F* (1, 16) = 10.03; *p*(treatment) = 0.0011 *F*(1, 16 = 15.72); **Fig. 7C**). Critically, however, cFos+ neuron numbers in CA1 were increased by ML297, even in SD mice (Sidak’s *post hoc* test vs. SD+Vehicle, *p* = 0.0058). This suggests that CA1 cFos+ cell numbers are enhanced by ML297 administration even in a scenario where consolidation of CFM has been disrupted by SD. Nonetheless, higher numbers of cFos+ neurons in CA1 was associated with better CFM recall (i.e., higher levels of freezing; Pearson correlation coefficient, *R* = 0.6701, *p* = 0.0012; **Fig. 7D**). cFos+ neuron numbers in CA3 showed a similar overall pattern, but varied significantly as a function of sleep only (two-way ANOVA; *p*(sleep condition) = 0.0030, *F* (1, 16) = 12.25; *p*(treatment) = 0.1914; *F*(1, 16 = 1.861); **Fig. 7C**). cFos+ cell counts in CA3 also reflected freezing levels during recall for individual animals (Pearson correlation coefficient, *R* = 0.5719, *p* = 0.0084; **Fig. 7D**).

**Figure 7:**
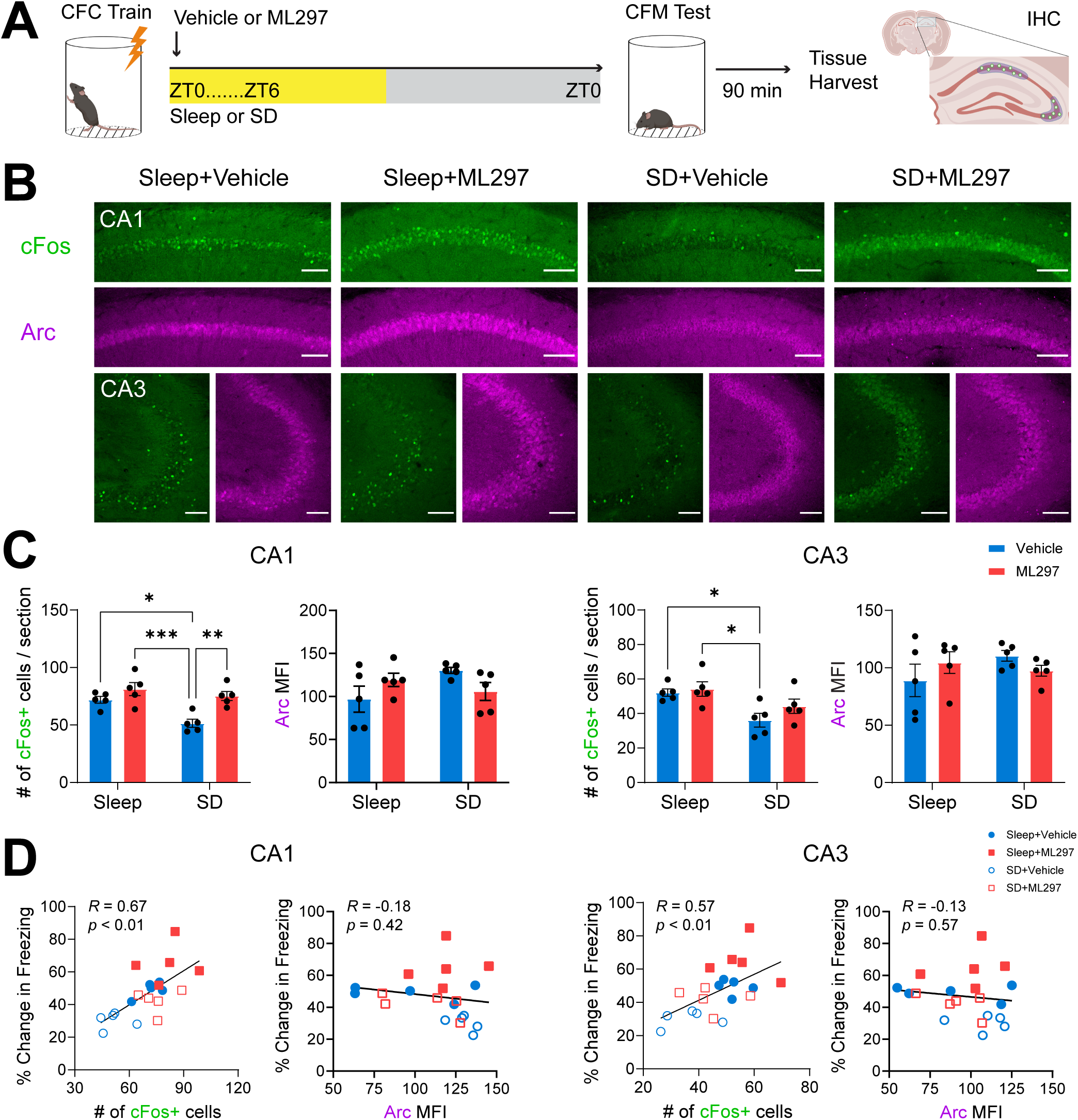
Post-CFC ML297 increases the number of active neurons in CA1 during subsequent CFM recall, in a sleep-independent manner. (*A*) Experimental design, as in Fig. 6A. (*B*) Representative images of cFos+ neurons (green) and Arc+ mean fluorescence intensity (MFI) in the pyramidal layers (magenta) in CA1 and CA3 following CFM recall in the four groups. Scale bar = 100 µm. (*C*) Post-CFC SD significantly decreased the number of cFos+ neurons in CA1 and CA3 during recall in vehicle-treated mice. In CA1 only, ML297 increased cFos+ neuron numbers after recall in SD mice as well as freely-sleeping mice. No significant changes in Arc MFI were observed with SD or drug treatment in either CA1 or CA3. *, **, and *** indicate *p* < 0.05, *p* < 0.01, and *p* < 0.001, respectively, Sidak’s *post hoc* test. (*D*) Numbers of cFos+ neurons in both CA1 and CA3 during recall correlated with CFM recall performance. No correlations were observed between freezing behavior and Arc MFI expression. *R* and *p* values are shown for Pearson correlation.

Due to the widespread nature of Arc expression in CA1 and CA3, we quantified mean fluorescence intensity (MFI) of Arc immunostaining in these sections after recall, using previous methods (12). Using this strategy for quantification of Arc, no significant differences were observed across groups for any of the groups in either CA1 or CA3 (**Fig. 7C**), and MFI values were not predictive of freezing behavior during recall (**Fig. 7D**).

Taken together, these studies suggest that both post-CFC sleep (vs. SD), and post-CFC administration of ML297, can increase dorsal hippocampus neuronal activation during subsequent CFM recall. These effects on hippocampal activation (particularly on neuronal activation in DG) during recall mirror, and are positively correlated with, freezing behavior during recall. These findings also support the idea that GIRK1 activation alters hippocampal network-level processes involved in consolidation in a sleep-dependent manner, leading to sleep-dependent changes in activation during recall.

## Discussion

We find that direct GIRK1 channel activation can increase post-CFC REM sleep, restoring REM sleep architecture (normally suppressed after fear learning) to baseline levels, and improving sleep-dependent consolidation of fear memory. We find that ML297-associated increases in overall REM sleep amounts, bout durations, and bout numbers (and reduced latency to REM sleep) co-occur with increased NREM spindle power over the first 6 h post-CFC. It is likely that these changes are driven by the same underlying mechanism, due to the increasingly well-established causal relationship between spindle-rich NREM sleep and transitions from NREM into REM sleep (61, 62). In other words, it is likely that transitions into REM sleep after ML297 administration reflect the normal physiology of such transitions. Critically, all of the NREM and REM sleep changes caused by ML297 in the hours following CFC appear to be renormalizing sleep architecture to what is typically observed under baseline conditions (i.e., in the absence of fear learning). Two features of these findings are worth noting. First, the ML297-induced changes in sleep architecture, which are correlated with successful CFM consolidation, all occur within a window of time (i.e., the first 6 h after CFC) where SD is sufficient to disrupt the consolidation process (6, 16–18). Second, post-CFC SD is sufficient to prevent the CFM consolidation benefits of ML297 administration.

It is worth noting that both spindle-rich NREM sleep (such as that present at the transition to REM) and REM sleep have been linked to memory storage, across species - from humans to rodent models (1, 64, 65). NREM spindles have received a great deal of recent study due to their linkage to sleep-related improvements on a range of mnemonic tasks, and to sleep-dependent synaptic plasticity in neocortex (3, 21, 33, 55, 66, 67). Within the hippocampus, spindles and other NREM-associated electrophysiological (19, 21) and neuromodulatory (11, 22) changes have been mechanistically linked to successful CFM consolidation. Our present data suggest that REM is at least equally vital for CFM, and support growing body of data indicating that REM-specific features of post-learning hippocampal activity (18, 28) and gene expression (68–70) are essential for the consolidation process. Together, our findings support the notion that post-CFC REM sleep plays a causal role in promoting fear memory consolidation.

To better understand the link between our behavioral results and hippocampal network-level events underlying successful memory consolidation, we examined IEG expression within the hippocampus following CFM recall. We find that just as recall itself is suppressed after post-CFC SD, the number of cFos+ and Arc+ neurons in DG after recall, as well as the number of cFos+ neurons in downstream regions CA3 and CA1, is decreased in SD mice. This is consistent with the idea that hippocampal activation during recall is reduced overall after SD-disrupted consolidation. Because mice are given an adequate opportunity for recovery sleep between SD and recall (i.e., 18 h from ZT6 to ZT0 the following day), we believe that this alteration is due to a long-term change in the strength of the memory trace itself, rather than an acute effect of SD on hippocampal activation (11, 12, 71). In other words, an increase in the number of neurons active at recall following *ad lib* sleep would reflect more neurons’ inclusion into the hippocampal “engram”, while decreases after SD would reflect a reduction in neuronal incorporation into the memory trace. Intriguingly, ML297 administration after CFC in freely-sleeping mice results in a further increase in the number of IEG+ neurons in DG, where recall-activated neurons are generally sparser, but does not affect IEG+ numbers when administered in the context of SD. These sleep-dependent effects of ML297 on DG neurons’ activity during recall closely reflects effects of ML297 on CFM consolidation. In contrast, in CA1, ML297 increases numbers of recall-activated cFos+ neurons, regardless of whether mice are freely-sleeping or sleep-deprived. This suggests additional, sleep-independent effects of ML297 within CA1, the region of hippocampus where GIRK1 channels are most abundant (44, 48, 52). Overall, our data suggest that GIRK channel activation has sleep-dependent and sleep-independent effects on the dorsal hippocampal network in the context of consolidation, leading to incorporation of more neurons into the CFM engram, which is evident in the pattern of network activation during CFM recall.

While numerous genetic findings suggest that loss of GIRK channel activity disrupts hippocampal memory processing (72), their precise molecular role in this process remains unclear. *In vitro* studies have shown that in the hippocampus (e.g. CA1) GIRK channel activation induces hyperpolarization, reduce neuronal excitability, and suppresses LTP (49, 50, 53). However, it is unknown how these effects translate to *in vivo* function, and particularly how these changes are modulated in different brain states (such as wake vs. NREM and REM sleep). Future studies will be needed to disentangle the relationship between direct cellular effects of ML297 administration, its behavioral effects (e.g., sleep-promoting, anxiolytic), and its mnemonic effects during the memory consolidation process.

Recent data have implicated GIRK channels as a target for therapeutics in various neurological and psychiatric conditions including epilepsy, Alzheimer’s disease, substance abuse, and anxiety disorders (52, 72). Our present data support the recent suggestion (40) that GIRK1 activation via ML297 could also be beneficial as a hypnotic. Beyond this, our data demonstrate that this hypnotic agent restores physiological REM sleep (whose disruption by fear learning is well-established (23)) to promote sleep-dependent memory consolidation. These findings have important ramifications for treatment of disorders - including neurodevelopmental disorders, dementia, and anxiety disorders) where both sleep architecture and cognitive function are disrupted.

## Acknowledgements

We thank the members of the Aton lab for useful feedback on these studies, and Gregg Sobocinski for microscopy support. This work was supported by a University of Michigan Rackham Graduate School Predoctoral Fellowship, Graduate Student Research Grant, and Merit Fellowship to JDM, a University of Michigan Kavli Neuroscience Innovators Magnificent Michigan Summer Fellowship to WPB, and NIH research grants R01 NS104776 and RF1 NS118440 to SJA.

## Supplementary Figure Legends

**Figure S1:**
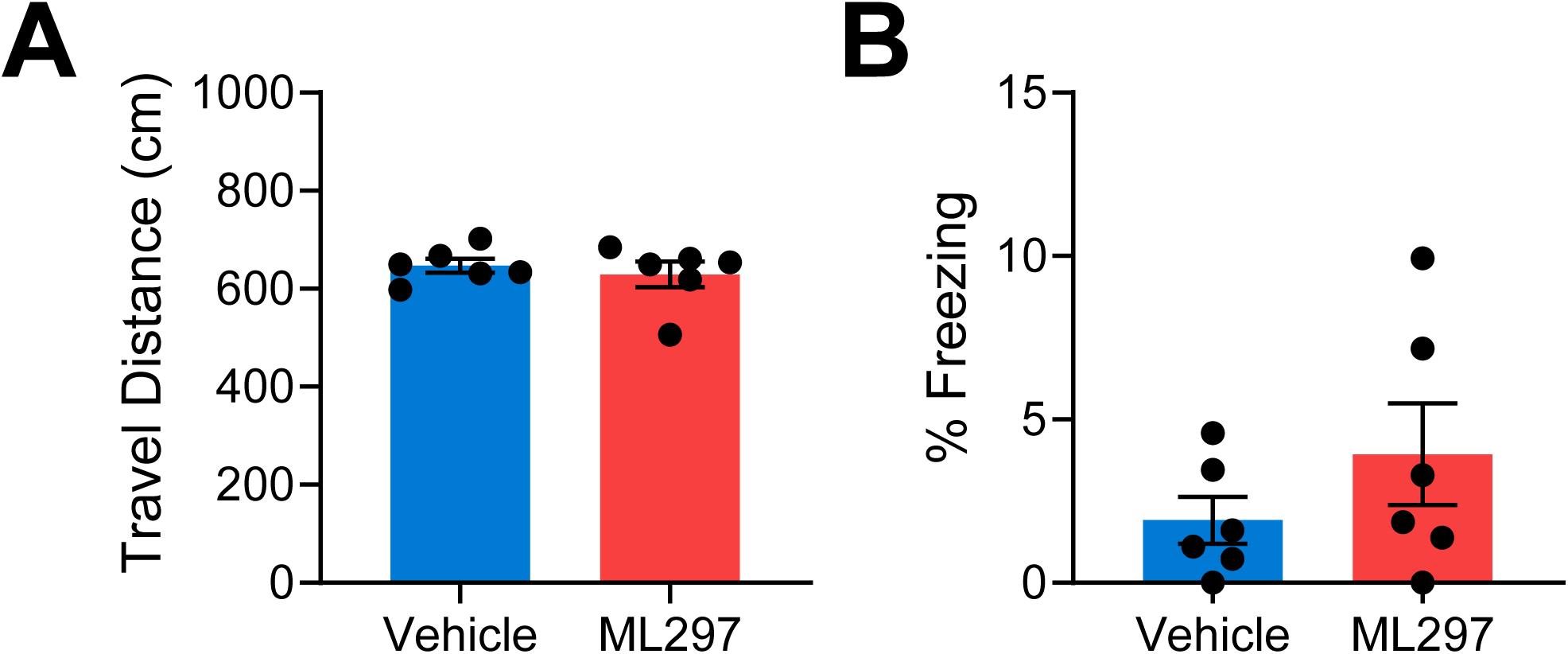
Travel distance and freezing levels are similar during initial training for EEG-implanted mice. Vehicle- and ML297-treated mice had similar travel distance (A) and freezing behavior (B) amounts during CFC (pre-shock). *n* = 6 mice/group

**Figure S2:**
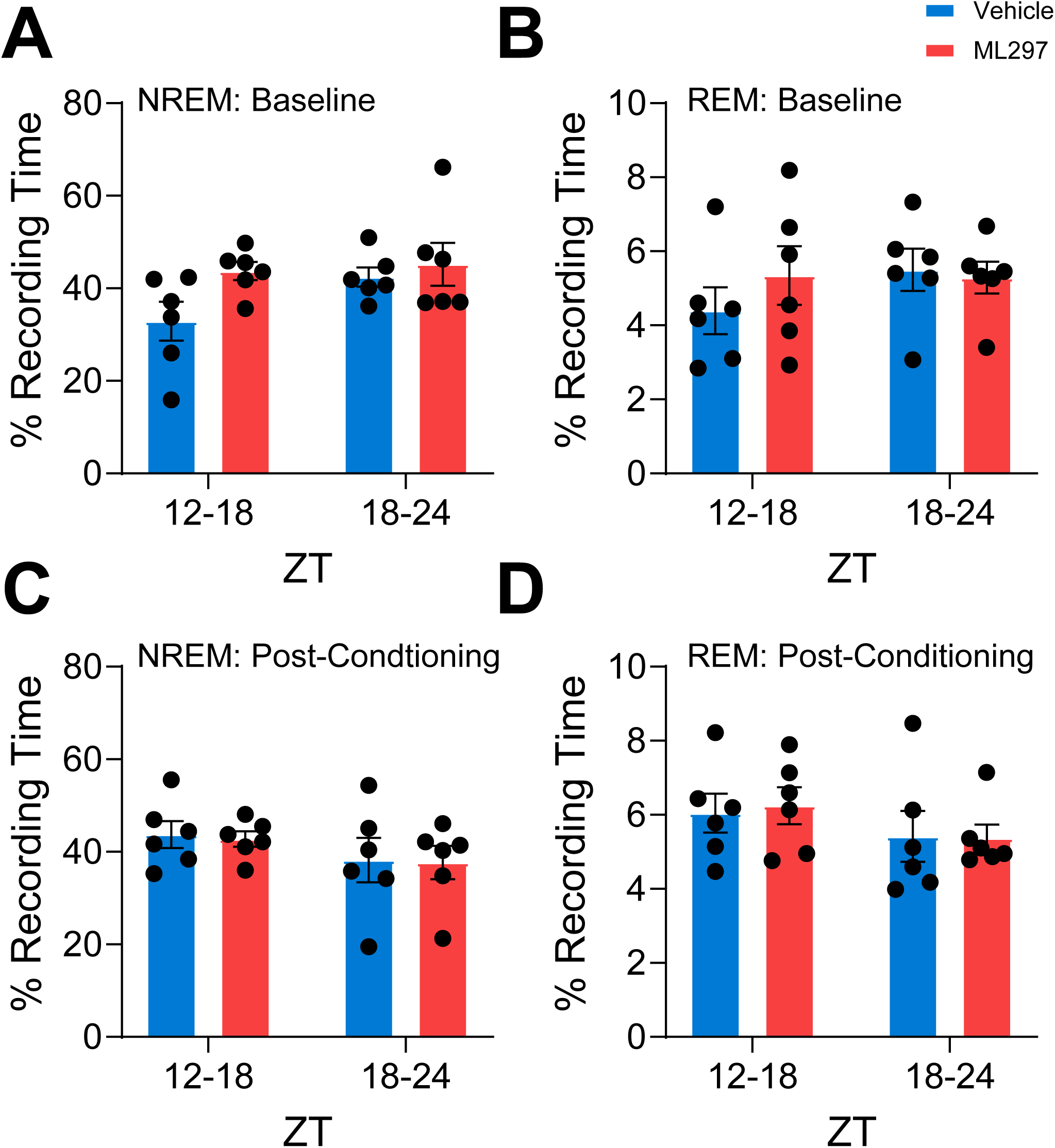
NREM and REM sleep architecture are similar during the dark cycle post-CFC. (***A***) NREM and (***B***) REM total sleep behavior across 6 h periods during dark cycle are similar for vehicle- and ML297-treated mice. Gray shaded areas represent lights off. *n* = 6 mice/group

**Figure S3:**
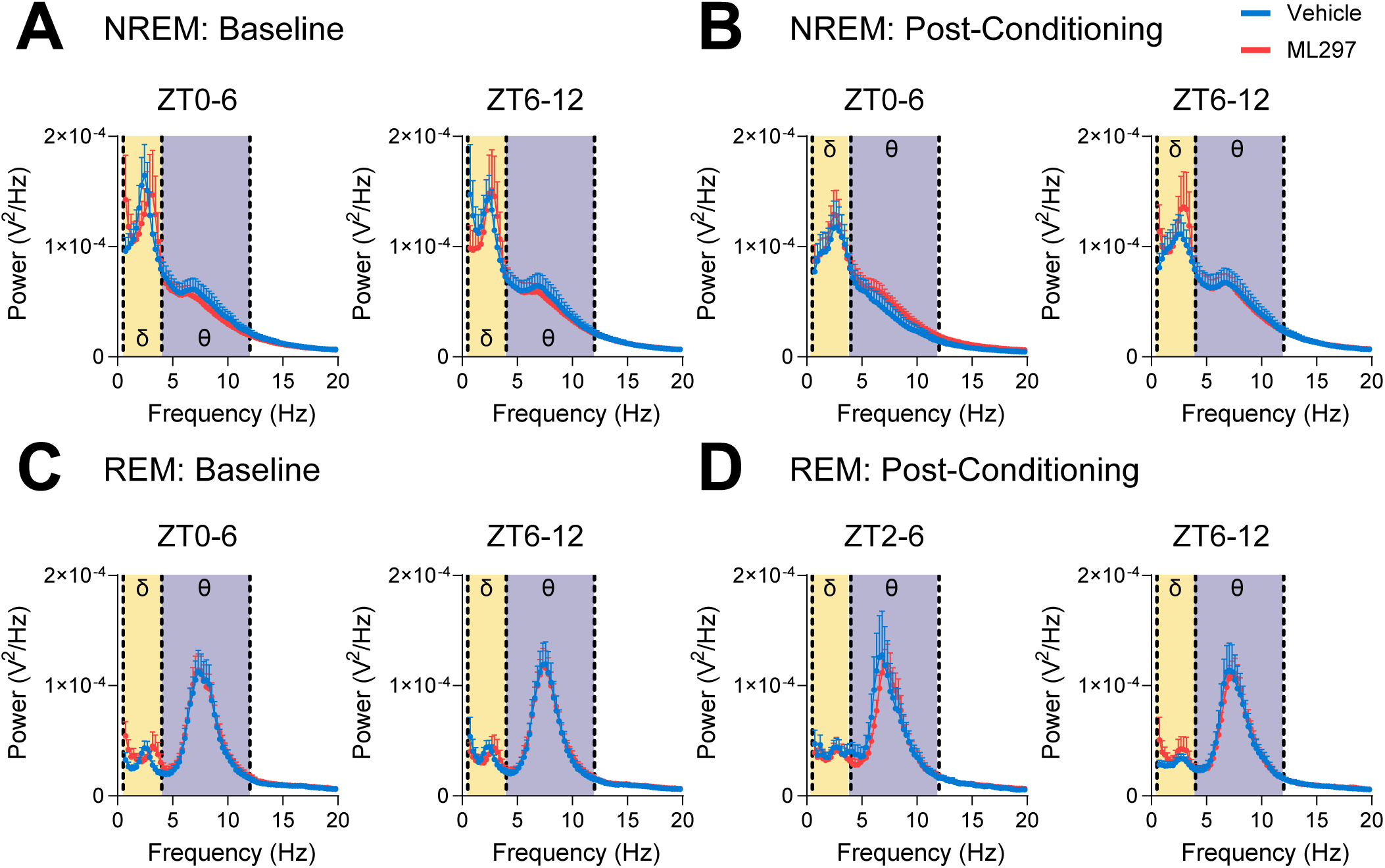
NREM and REM raw EEG spectral power. EEG raw power spectra (recorded over visual cortex, bilaterally) are shown for vehicle- and ML297-treated mice during NREM baseline (***A***) and post-CFC (***B***), and during REM baseline (***C***) and post-CFC (***D***). Values indicate raw voltage of spectral power at each frequency band, *n* = 6 mice/group.

**Figure S4:**
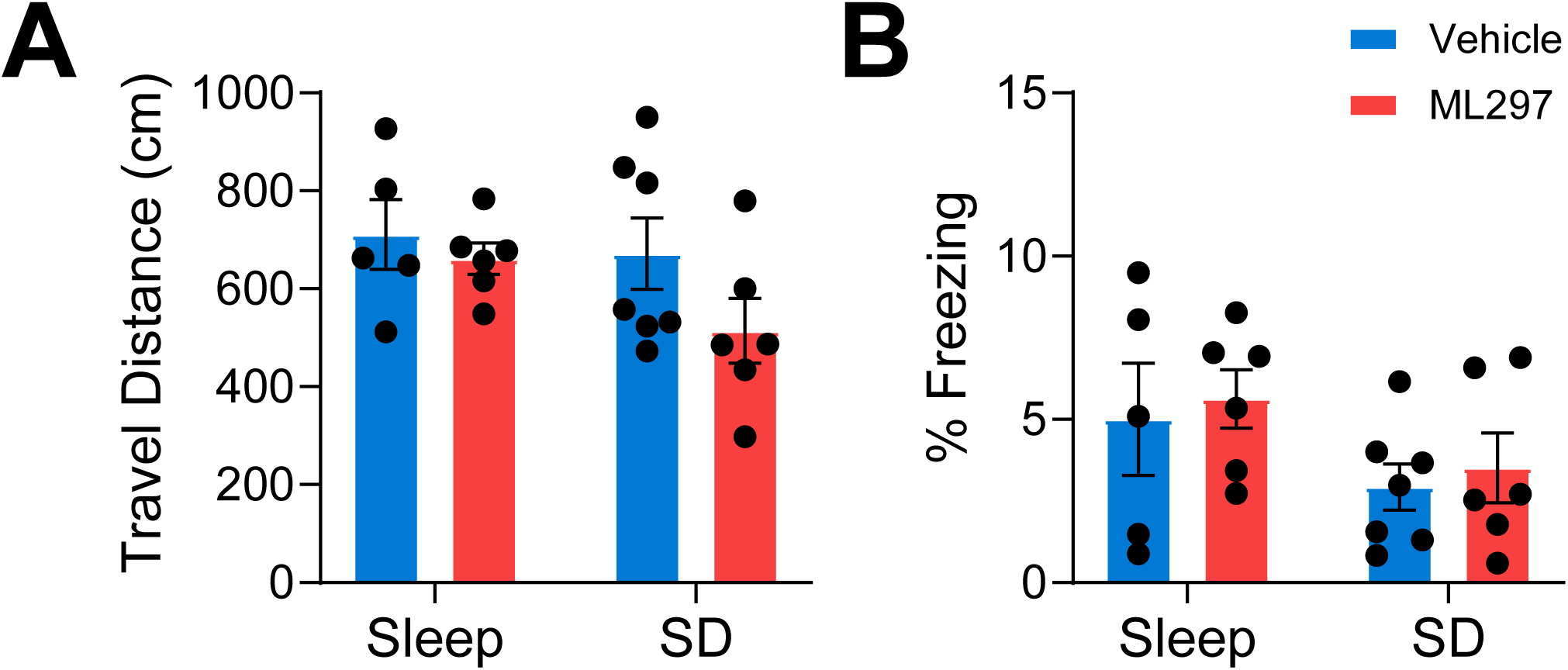
Travel distance and freezing levels are similar during initial training for non-EEG-implanted mice. Vehicle- and ML297-treated mice for both *ad lib* sleep and sleep deprivation groups had similar travel distance (***A***) and freezing behavior (***B***) amounts during training (pre-shock). *n* = 5 and 6 mice, respectively, for vehicle and ML297 with *ad lib* sleep, and *n* = 7 and 6 mice, respectively, for vehicle and ML297 with SD.

## References

1. B. Rasch, J. Born, About sleep’s role in memory. Physiol Rev 93, 681–766 (2013).

2. R. Stickgold, Parsing the role of sleep in memory processing. Curr Opin Neurobiol 23, 847–853 (2013).

3. C. Puentes-Mestril, J. Roach, N. Niethard, M. Zochowski, S. J. Aton, How rhythms of the sleeping brain tune memory and synaptic plasticity. Sleep 42, pii: zsz095 (2019).

4. C. Puentes-Mestril, S. J. Aton, Linking network activity to synaptic plasticity during sleep: hypotheses and recent data. Frontiers in Neural Circuits 11, doi: 10.3389/fncir.2017.00061 (2017).

5. C. Puentes-Mestril et al., Sleep loss drives brain region- and cell type-specific alterations in ribosome-associated transcripts involved in synaptic plasticity and cellular timekeeping. J Neurosci 41, 5386–5398 (2021).

6. J. Delorme et al., Hippocampal neurons’ cytosolic and membrane-bound ribosomal transcript profiles are differentially regulated by learning and subsequent sleep. Proc Natl Acad Sci USA 118 (2021).

7. S. B. Noya et al., The Forebrain Synaptic Transcriptome Is Organized by Clocks but Its Proteome Is Driven by Sleep Science 366 (2019).

8. F. Bruning et al., Sleep-wake Cycles Drive Daily Dynamics of Synaptic Phosphorylation Science 366 (2019).

9. J. G. Klinzing, N. Niethard, J. Born, Mechanisms of systems memory consolidation during sleep. Nature Neuroscience 22, 1598–1610 (2019).

10. C. N. Hor et al., Sleep-wake-driven and circadian contributions to daily rhythms in gene expression and chromatin accessibility in the murine cortex. Proc Natl Acad Sci USA 116, 25773–25783 (2019).

11. J. Delorme et al., Sleep loss drives acetylcholine- and somatostatin interneuron-mediated gating of hippocampal activity, to inhibit memory consolidation. Proc Natl Acad Sci USA 118 (2021).

12. J. E. Delorme, V. Kodoth, S. J. Aton, Sleep loss disrupts Arc expression in dentate gyrus neurons. Neurobiol Learn Mem 160, 73–82 (2019).

13. R. Havekes, S. J. Aton, Impacts of Sleep Loss Versus Waking Experience on Brain Plasticity: Parallel or Orthogonal? Trends in Neuroscience 43, 385–393 (2020).

14. R. Havekes, T. Abel, The tired hippocampus: the molecular impact of sleep deprivation on hippocampal function. Curr Opin Neurobiol 44, 13–19 (2017).

15. T. M. Prince et al., Sleep deprivation during a specific 3-hour time window post-training impairs hippocampal synaptic plasticity and memory. Neurobiol Learn Mem 109, 122–130 (2014).

16. L. A. Graves, E. A. Heller, A. I. Pack, T. Abel, Sleep deprivation selectively impairs memory consolidation for contextual fear conditioning. Learn. Mem. 10, 168–176 (2003).

17. C. G. Vecsey et al., Sleep deprivation impairs cAMP signalling in the hippocampus. Nature 461, 1122–1125 (2009).

18. N. Ognjanovski, C. Broussard, M. Zochowski, S. J. Aton, Hippocampal Network Oscillations Rescue Memory Consolidation Deficits Caused by Sleep Loss. Cereb. Cortex 28, 3711–3723 (2018).

19. N. Ognjanovski et al., Parvalbumin-expressing interneurons coordinate hippocampal network dynamics required for memory consolidation. Nature Communications 8, 15039 (2017).

20. N. Ognjanovski, D. Maruyama, N. Lashner, M. Zochowski, S. J. Aton, CA1 hippocampal network activity changes during sleep-dependent memory consolidation. Front Syst Neurosci 8, 61 (2014).

21. F. Xia et al., Parvalbumin-positive interneurons mediate neocortical-hippocampal interactions that are necessary for memory consolidation. eLife 6, e27868 (2017).

22. Q. M. Skilling et al., Acetylcholine-gated current translates wake neuronal firing rate information into a spike timing-based code in Non-REM sleep, stabilizing neural network dynamics during memory consolidation PLoS Computational Biology 17, e1009424 (2021).

23. X. Tang, L. Yang, L. D. Sanford, Interactions between brief restraint, novelty and footshock stress on subsequent sleep and EEG power in rats. Brain Research In Press, Corrected Proof (2007).

24. L. Sanford, A. J. Silvestri, R. J. Ross, A. R. Morrison, Influence of fear conditioning on elicited ponto-geniculate-occipital waves and rapid eye movement sleep. Arch Ital Biol 139, 169–183 (2001).

25. M. Menz et al., The role of sleep and sleep deprivation in consolidating fear memories Neuroimage 75, 87–96 (2013).

26. V. Spoormaker, G. Gvozdanovic, P. Samann, M. Czisch, Ventromedial prefrontal cortex activity and rapid eye movement sleep are associated with subsequent fear expression in human subjects. Experimental Brain Res 232, 1547–1554 (2014).

27. E. Pace-Schott, A. Germain, M. Milad, Effects of sleep on memory for conditioned fear and fear extinction. Psychol Bull 141, 835–857 (2015).

28. R. Boyce, S. D. Glasgow, S. Williams, A. Adamantidis, Causal evidence for the role of REM sleep theta rhythm in contextual memory consolidation. Science 352, 812–816 (2016).

29. S. J. Aton et al., The sedating antidepressant trazodone impairs sleep-dependent cortical plasticity. PLoS ONE 4, e6078 (2009).

30. J. Seibt et al., The non-benzodiazepine hypnotic Zolpidem impairs sleep-dependent cortical plasticity. SLEEP In Press (2008).

31. J. Hall-Porter, P. Schweitzer, R. Eisenstein, H. Ahmed, J. Walsh, The Effect of Two Benzodiazepine Receptor Agonist Hypnotics on Sleep-Dependent Memory Consolidation. J Clin Sleep Med 10, 27–34 (2014).

32. E. J. Wamsley et al., The effects of eszopiclone on sleep spindles and memory consolidation in schizophrenia: a randomized placebo-controlled trial. Sleep 36, 1369–1376 (2013).

33. S. C. Mednick et al., The critical role of sleep spindles in hippocampal-dependent memory: a pharmacology study. J Neurosci 33, 4494–4504 (2013).

34. J. Vienne et al., Differential effects of sodium oxybate and baclofen on EEG, sleep, neurobehavioral performance, and memory. Sleep 35, 1071–1083 (2012).

35. C. Leong et al., A systematic scoping review of the effects of central nervous system active drugs on sleep spindles and sleep-dependent memory consolidation. Sleep Med Rev 62, 101605 (2022).

36. L. P. Carter, D. Pardi, J. Gorsline, R. S. Griffiths, Illicit gamma-hydroxybutyrate (GHB) and pharmaceutical sodium oxybate (Xyrem®): Differences in characteristics and misuse. Drug Alcohol Depend 104, 1–10 (2009).

37. O. C. Snead, Evidence for GABAB-mediated mechanisms in experimental generalized absence seizures. Eur J Pharmacol 213, 342–349 (1992).

38. T. Roth et al., Effect of sodium oxybate on disrupted nighttime sleep in patients with narcolepsy. . J Sleep Res 26, 407–417 (2017).

39. J. Black, D. Pardi, C. S. Hornfeldt, N. Inhaber, The nightly administration of sodium oxybate results in significant reduction in the nocturnal sleep disruption of patients with narcolepsy. . Sleep Med 10, 829–835 (2009).

40. B. Zou et al., Direct activation of G-protein-gated inward rectifying K+ channels promotes nonrapid eye movement sleep. Sleep 42 (2019).

41. E. Szabadi, Selective targets for arousal-modifying drugs: implications for the treatment of sleep disorders Drug Discov Today 19, 701–708 (2014).

42. A. Kuriyama, M. Honda, Y. Hayashino, Ramelteon for the treatment of insomnia in adults: a systematic review and meta-analysis. Sleep Med 15, 385–392 (2014).

43. A. Inanobe et al., Characterization of G-protein-gated K+ channels composed of Kir3.2 subunits in dopaminergic neurons of the substantia nigra. J Neurosci 19, 1006–1101 (1999).

44. A. Tinker, Y. N. Jan, L. Y. Jan, Regions responsible for the assembly of inwardly rectifying potassium channels. Cell 87, 858–868 (1996).

45. K. B. Walsh, Targeting GIRK channels for the development of new therapeutic agents. Front Pharmacol 2, 64 (2011).

46. D. E. Logothetis et al., Unifying Mechanism of Controlling Kir3 Channel Activity by G Proteins and Phosphoinositides. Int Rev Neurobiol 123, 1–26 (2015).

47. S. D. Carter, K. R. Mifsud, J. M. Reul, Distinct epigenetic and gene expression changes in rat hippocampal neurons after Morris water maze training. Front Behav Neurosci 9, 156 (2015).

48. V. Kloukina et al., G-protein-gated inwardly rectifying K+ channel 4 (GIRK4) immunoreactivity in chemically defined neurons of the hypothalamic arcuate nucleus that control body weight. J Chem Neuroanat 44, 14–23 (2012).

49. C. Luscher, L. Y. Jan, M. Stoffel, R. C. Malenka, R. A. Nicoll, G protein-coupled inwardly rectifying K+ channels (GIRKs) mediate postsynaptic but not presynaptic transmitter actions in hippocampal neurons. . Neuron 19, 687–695 (1997).

50. S. Signorini, Y. J. Liao, S. A. Duncan, L. Y. Jan, M. Stoffel, Normal cerebellar development but susceptibility to seizures in mice lacking G protein-coupled, inwardly rectifying K+ channel GIRK2. . Proc Natl Acad Sci USA 94, 923–927 (1997).

51. N. Wydeven et al., Mechanisms underlying the activation of G-protein–gated inwardly rectifying K+ (GIRK) channels by the novel anxiolytic drug, ML297. Proc Natl Acad Sci USA 111, 10755–10760 (2014).

52. K. Kaufmann et al., ML297 (VU0456810), the First Potent and Selective Activator of the GIRK Potassium Channel, Displays Antiepileptic Properties in Mice. ACS Chem Neurosci 4, 1278–1286 (2013).

53. S. Djebari et al., G-Protein-Gated Inwardly Rectifying Potassium (Kir3/GIRK) Channels Govern Synaptic Plasticity That Supports Hippocampal-Dependent Cognitive Functions in Male Mice. J Neurosci 41, 7086–7102 (2021).

54. B. C. Clawson et al., Causal role for sleep-dependent reactivation of learning-activated sensory ensembles for fear memory consolidation. Nat Communications 12 (2021).

55. J. Durkin et al., Cortically coordinated NREM thalamocortical oscillations play an essential, instructive role in visual system plasticity. Proceedings National Academy of Sciences 114, 10485–10490 (2017).

56. A. C. Pawlyk, A. R. Morrison, R. J. Ross, F. X. Brennan, Stress-Induced Changes in Sleep in Rodents: Models and Mechanisms. Neurosci Biobehav Rev 32, 99–117 (2008).

57. S. Uchida, T. Maloney, J. D. March, R. Azari, I. Feinberg, Sigma (12–15 Hz) and delta (0.3-3 Hz) EEG oscillate reciprocally within NREM sleep. . Brain Res Bull 27, 93–96 (1991).

58. L. Trachsel, I. Tobler, A. A. Borbely, Electroencephalogram analysis of non-rapid eye movement sleep in rats. . Am J Physiol 255, R27–37 (1988).

59. S. Sarasso et al., Thalamic and neocortical differences in the relationship between the time course of delta and sigma power during NREM sleep in humans. J Sleep Res 30, e13166 (2021).

60. C.-F. V. Latchoumane, H.-V. V. Ngo, J. Born, H. S. Shin, Thalamic Spindles Promote Memory Formation during Sleep through Triple Phase-Locking of Cortical, Thalamic, and Hippocampal Rhythms. Neuron 95, 424–435 (2017).

61. M. Bandarabadi et al., A role for spindles in the onset of rapid eye movement sleep Nat Communications 11 (2020).

62. A. Kim et al., Optogenetically induced sleep spindle rhythms alter sleep architectures in mice. Proc Natl Acad Sci U S A 109, 20673–20678 (2012).

63. X. Liu et al., Optogenetic stimulation of a hippocampal engram activates fear memory recall. Nature 484, 381–385 (2012).

64. S. Ribeiro, M. A. L. Nicolelis, Reverberation, storage, and postsynaptic propagation of memories during sleep. Learn.Mem. 11, 686–696 (2004).

65. R. Stickgold, M. P. Walker, Sleep-dependent memory consolidation and reconsolidaton. Sleep Med 8, 331–343 (2007).

66. B. C. Clawson, J. Durkin, S. J. Aton, Form and function of sleep spindles across the lifespan. Neural Plast. 2016, 6936381 (2016).

67. S. J. Aton et al., Visual experience and subsequent sleep induce sequential plastic changes in putative inhibitory and excitatory cortical neurons. Proc Natl Acad Sci U S A 110, 3101–3106 (2013).

68. J. B. Calais, E. B. Ojopi, E. Morya, K. Sameshima, S. Ribeiro, Experience-dependent upregulation of multiple plasticity factors in the hippocampus during early REM sleep. Neurobiol Learn Mem 122, 19–27 (2015).

69. S. Datta, G. Li, S. Auerbach, Activation of phasic pontine-wave generator in the rat: a mechanism for expression of plasticity-related genes and proteins in the dorsal hippocampus and amygdala. Eur J Neurosci 27, 1876–1892 (2008).

70. J. Ulloor, S. Datta, Spatio-temporal activation of cyclic AMP response element-binding protein, activity-regulated cytoskeletal-associated protein and brain-derived nerve growth factor: a mechanism for pontine-wave generator activation-dependent two-way active-avoidance memory processing in the rat. Journal of Neurochemistry 95, 418–428 (2005).

71. S. S. Yoo, P. T. Hu, N. Gujar, F. A. Jolesz, M. P. Walker, A deficit in the ability to form new human memories without sleep. Nat Neurosci (2007).

72. J. Mayfield, Y. A. Blednov, R. A. Harris, Behavioral and Genetic Evidence for GIRK Channels in the CNS: Role in Physiology, Pathophysiology, and Drug Addiction. Int Rev Neurobiol 123, 279–313 (2015).

